# Prior Signal Acquisition Software Versions for Orbitrap Underestimate Low Isobaric Mass Tag Intensities, Without Detriment to Differential Abundance Experiments

**DOI:** 10.1101/2021.07.18.452841

**Authors:** Tom S Smith, Anna Andrejeva, Josie A. Christopher, Oliver M. Crook, Mohamed A. W. Elzek, Kathryn S Lilley

## Abstract

Tandem mass tags (TMT) enable simple and accurate quantitative proteomics for multiplexed samples by relative quantification of tag reporter ions. Orbitrap™ quantification of reporter ions has been associated with a characteristic notch region in intensity distribution, within which few reporter intensities are recorded. This has been resolved in version 3 of the instrument acquisition software, Tune. However, 53 % of Orbitrap Fusion, Lumos or Eclipse submissions to PRIDE were generated using prior software versions. To quantify the impact of the notch on existing quantitative proteomics data, we generated a mixed species benchmark and acquired quantitative data using Tune versions 2 and 3. Sub-notch intensities are systemically underestimated with Tune version 2, leading to over-estimation of the true differences in intensities between samples. However, when summarising reporter ion intensities to higher level features, such as peptides and proteins, few features are significantly affected. Targeted removal of spectra with reporter ion intensities below the notch is not beneficial for differential peptide or protein testing. Overall, we find the systematic quantification bias associated with the notch is not detrimental for a typical proteomics experiment.

## Introduction

Bottom-up quantitative proteomics entails proteolytic digestion of proteins to peptides, quantification of peptide abundances by mass spectrometry (MS) and summarisation of protein abundances. Data-dependent acquisition (DDA) is the classical acquisition mode. Due to the stochastic nature of peptide selection, not all peptides present in a sample are fragmented and sequenced, with data completeness diminishing as the number of samples increases ^1^. Missing values are reduced with data-independent acquisition (DIA) approaches ^2^, although there are still detection threshold limits and samples cannot be multiplexed in typical DIA workflows.

Alternatively, samples may be labelled with isobaric tags which possess the same mass, but different distributions of heavy isotopes within the tags, such that a sample-specific mass reporter tag is released by high energy collision-induced dissociation ^3^. By enabling sample multiplexing, the frequency of missing values is thus reduced compared to Label Free Quantification ^4^. Furthermore, all samples are quantified from the same peptide spectrum matched (PSM) ions, greatly simplifying summarisation to protein-level abundances ^5^.

Tandem Mass Tags (TMT) are the most commonly used isobaric tags, with current chemistry allowing up to 18 samples to be multiplexed ^6^. Since quantification is typically of the tag rather than the peptide, TMT proteomics has been shown to suffer ratio compression by the presence of co-isolated ‘interference’ peptides ^7^. Such compression can be partially mitigated by use of synchronous precursor selection (SPS)-MS3 quantification ^8^. Thus, robust quantification of TMT labelled peptides requires high-resolution, accurate mass spectrometers, with Orbitrap™ devices being commonly employed. Intriguingly, a recent characterisation of TMT reporter ion signals obtained from Orbitrap identified a consistent absence of intensities within a specific range, which visually appears as ‘notch’ in the distribution of intensities ^9^. The presence of the notch was determined to depend on the automatic gain control (AGC) target and maximum injection time Orbitrap parameters, with higher values reducing its prominence. Hughes *et al* hypothesised the cause of the notch is rooted in the signal processing behaviour, from their observation of a notch in all datasets acquired via Orbitrap-based measurements, regardless of the other associated MS hardware, and the exhaustive consideration of user-defined parameters. Hughes *et al* further speculated that standard procedures to remove low-intensity spectra would mitigate any potential issues with quantification inaccuracies.

The release notes for the Orbitrap Fusion Series 3.0 signal acquisition software, Tune, explain that the notch has been resolved in Item DE 54684: ‘Addressed the peak intensity (linearity) for extremely low S/N values’ ^10^. Nevertheless, published datasets have used prior software versions and trust in the results from these experiments is contingent upon accurate quantification.

Here, we further examine the ‘notch’ phenomenon and demonstrate what happens to reporter ion intensities that fall inside the notch. Crucially, we examine the overall impact of the notch for the detection of significant fold changes and estimation of their magnitude, in an experiment that aims to measure differential abundance.

## Results and discussion

To estimate the proportion of published Orbitrap datasets using Tune versions that will generate a notch, we downloaded .raw files from PRIDE for all studies using only Orbitrap Fusion, Lumos or Eclipse hardware since these use the same series of acquisition software. We then extracted the version of Tune used from the .raw file (see methods). In total, 1283 studies were examined, with the remaining studies either falsely stating the instrument model, or not possessing files that could be parsed for meta data (Figure 1a). Overall, 53.0 % of studies used Tune versions that generate a notch, including 27.3 % of submissions in the year to 28^th^ June 2021. It is therefore imperative to determine what impact the notch has on quantitative proteomics.

**Figure 1.**
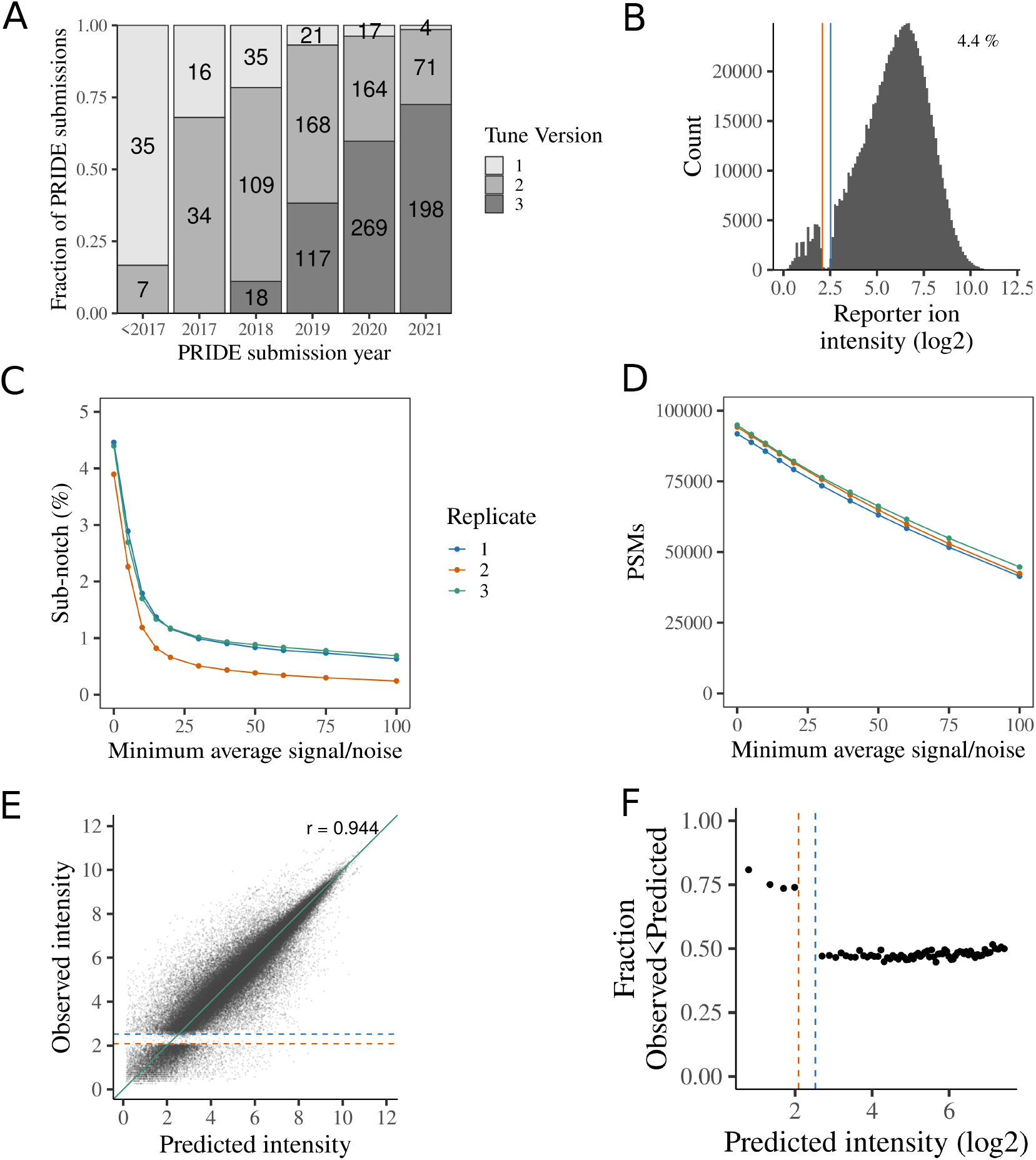
(A) Tune major versions for Orbitrap Lumos, Fusion and Eclipse PRIDE submissions (B) The distribution of ion signals for TMT reporters U-2 OS LOPIT-DC, replicate 1. The percentage of tag intensities below the upper boundary of the notch is stated in the top right corner. (C-D) The impact of filtering PSMs by their average signal/noise. (C) The proportion of reporter tag intensities at or below the notch. (D) The number of PSMs remaining. (E) Observed tag intensities vs predicted tag intensities. Green line is equality. Pearson product-moment correlation coefficient shown in the top right corner (F) Fraction of observed tag intensities that are below the prediction for binned predicted tag intensity. Observed tag intensities below the notch are systematically underestimated relative to the prediction. The approximate boundaries of the notch region are denoted by vertical or horizontal lines in B, E and F. Equivalent plots for replicate 2 and 3 are shown in Figure S1–2.

We first reanalysed previously published Orbitrap SPS-MS3 TMT data acquired using Tune v2 to demonstrate that average signal/noise filtering is not a sufficient remedy for the notch. Between 3.9-4.5 % tag intensities fell within or below the notch (Figure 1b & S1b-c) in our U-2 OS LOPIT-DC ^11^ experiments, with the proportion varying considerably between tags within a given experiment (Figure S1d-f). Using increasingly stringent filtering to remove PSMs with low average signal/noise ratios reduces the proportion of sub-notch values (Figure 1c-d). However, even after removing PSMs with average signal/noise less than 100, 0.24-0.69 % of remaining tag intensities are below the notch, whilst 52.8-55.0 % of quantified PSMs are discarded. Using a more moderate 10-fold filter, as previously suggested ^12^, removes 6.6-6.8 % of the quantified PSMs, with 1.2-1.8 % of the remaining reporter intensities being below the notch. Thus, while average signal/noise filtering will increase quantification accuracy, it is not a complete remedy for sub-notch values.

To demonstrate what happens to notch intensities, we used redundant PSMs to predict expected reporter ion intensities. In brief, we considered sets of PSMs from the same peptide sequence and used the reporter ion intensities from the highest intensity PSM to predict the expected reporter ion intensity values for the other PSMs (see methods). Predicted and observed tag intensities were highly correlated, confirming the validity of this approach (Figure 1e, S2a-b). As expected, intensities below the notch are systematically under-estimated relative to the prediction (Figure 1f, S2c-d).

To explore the impact of the notch and sub-notch region on a typical quantitative TMT MS3 proteomics experiment, we considered suitable published benchmark experiments. Hughes *et al* performed a spike-in benchmark experiment to assess the impact of the notch ^9^. However, this included only 550 spike-in peptides, precluding a consideration of the impact of the notch in a routine proteomics experiment. In another benchmarking study, O’Connell *et al* spiked peptides from *S.cerevisiae* cell lysates into *H.sapiens* peptide samples at known concentration, thus mimicking a control vs treatment(s) differential protein abundance experimental design ^4^. However, the authors used a relatively high AGC target of 20,000 and maximum injection time of 150 ms. Thus, only 0.6 % tag intensities were observed at or below the notch, preventing any analysis of the impact of the notch in a more typical setting (Figure S3a). We therefore created our own spike-in benchmark experiment, aiming to observe a notch with typical prominence (see methods). Proteins extracted from whole cell lysates of *S.cerevisiae* and *H.sapiens* osteosarcoma cell line U-2 OS were digested to peptides and mixed, such that *S.cerevisiae* peptides were at 1x (5μg), 2x and 6x volumes in a total of 100 μg peptide. Thus, we generated differences in protein abundance between 1.06 - 6-fold for *H.sapiens* and *S.cerevisiae* proteins. Peptides were then labeled with TMT and quantified on an Orbitrap Mass Spectrometer with Tune 2.1 and Tune 3.4, with an AGC target of 50,000 and max injection time of 120 ms, which we expected to yield a moderate and typical notch ^9^ (see methods for details). Using these parameters and Tune 2.1, 3.2 % of tag intensities were within or below the notch (Figure 2b), with *S.cerevisiae* peptides showing a slightly greater proportion of low tag intensities (Figure S3b-c). In total we obtained 109,779 PSMs from 10,982 protein groups, of which 8,335 and 2,469 could be assigned to *H.sapiens and S. cerevisiae*, respectively.

**Figure 2.**
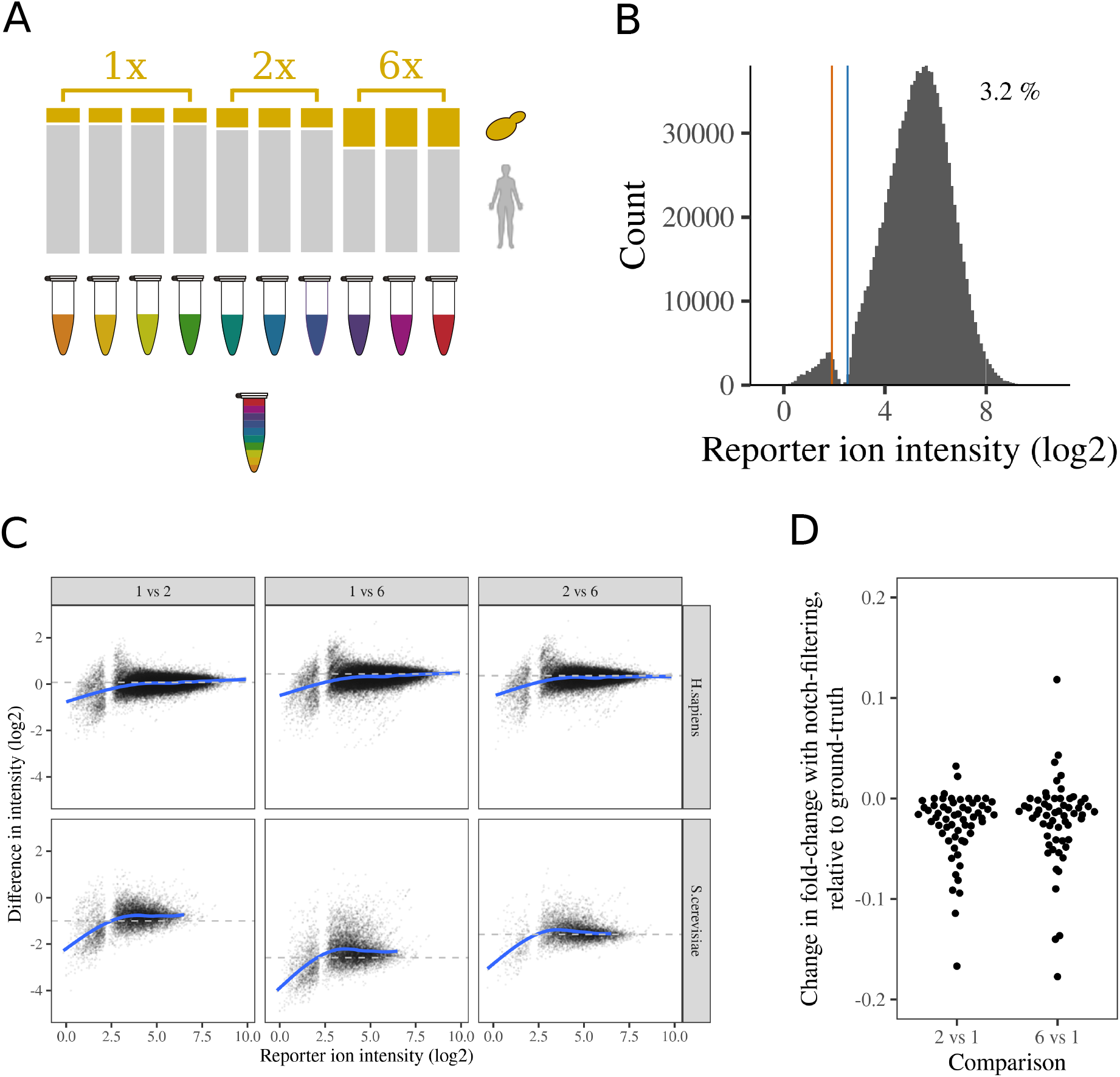
Impact of the notch on a differential expression benchmark proteomics experiment. (A) Schematic representation of benchmark TMT proteomics experimental design. Yeast peptides were spiked into human peptides at 1x, 2x and 6x volumes to induce ground truth fold-changes for both yeast and human proteins and labelled with TMT (B) The distribution of reporter ion intensities for TMT reporters. The approximate boundaries of the notch region are denoted by vertical lines. The percentage of tag intensities below the upper boundary of the notch is stated in the top right corner. (C) The difference between a single tag intensity and the mean tag intensity for a comparator group of tags. The ground truth is denoted by a dashed horizontal line. The blue line presents a generalized additive smoothing model for the relationship between tag intensity and intensity difference. (D) The difference in fold change when including notch filtering, relative to the ground truth. Positive values represent fold-change estimates closer to ground truth upon notch filtering. Only yeast proteins with at least one sub-notch reporter ion intensity are shown.

We applied strict filtering to minimise the possibility of PSM mis-identification or interference, demanding co-isolation < 10 %, and delta score > 0.5, with the later ensuring that rank 2 peptide matches for a given spectrum have a score less than half the rank 1 peptide match. We further removed PSMs with average reporter signal/noise < 10. Using these thresholds, 19,975 PSMs were retained. As expected, sub-notch tag intensities were clearly underestimated, with the correct fold-change only observed when reporter ion intensities were above the notch (Figure 2c). In contrast, using Tune 3.4, no notch was visible (Figure S4a) and the underestimation of fold-changes at low reporter ion intensity was largely resolved (Figure S4b).

We then aggregated reporter intensities to protein-level intensities, requiring at least two PSMs per protein. The majority of protein-level quantification estimates did not involve sub-notch intensities, with a maximum of 63 proteins (1.7%) having 25 % or greater sub-notch ion intensities in any given tag (Figure S5a).

To formally test the impact of the notch on the detection of peptide or protein differential abundance, we used *limma* to detect differential abundance, with and without prior filtering to remove PSMs containing intensities at or below the notch. The accuracy for identification of changes in intensity was measured by the F1 score, the harmonic mean of precision and recall. Fold-change estimates were very slightly closer to ground truth without notch filtering, but the difference was negligible (Figure S5b). For example, median fold-changes for *S.cerevisiae* proteins in the 6x vs 1x comparison decreased from 4.963 to 4.949 (Figure S6a). Crucially, notch filtering did not improve the F1 score for the most difficult to identify changes (*H.sapiens* 2x vs 1x) which was 0.245 and 0.246, respectively, with and without notch filtering (Figure S6b). Similarly, for the easier to detect fold changes, F1 was not significantly affected by notch filtering.

To ensure that our observations were not dependent on the PSM filtering, we varied the thresholds for maximum interference (10%, 50%, 100%), minimum average signal/noise (0, 10) and minimum Delta score (0, 0.2, 0.5). Each combination of PSM filtering thresholds was repeated with and without removing PSMs with sub-notch values. Thus, in total, 36 PSM filtering regimes were compared, representing a comprehensive set of PSM filtering schemas. Regardless of the thresholds used, notch filtering had negligible impact on fold-change estimates or the F1 score, whilst always reducing the number of proteins that could be interrogated (Figure S6). Thus, removing PSMs with sub-notch values does not appear to be beneficial and may even reduce sensitivity in differential protein abundance experiments.

Finally, we repeated the differential abundance testing at the peptide-level. Whilst differential abundance testing is more typically performed at the protein level, there are experiments where peptide-level testing is more appropriate, including in the analysis of post-translational modifications^12^. Given fewer reporter ion intensities are used to quantify peptides, we expected a more significant impact from the removal of sub-notch intensities. However, no improvements in the fold-changes were observed (Figure S5c) and the increase in F1 score was typically 0.001-0.005 and thus too slight to justify notch filtering (Figure S7).

To inspect the impact of the notch filtering on the estimated fold changes more directly, we considered proteins where at least one PSM was removed in the filtering, and compared the fold changes with and without notch filtering. As previously described, the proportion of proteins with a sub-notch PSM is relatively low. When using stringent PSM filtering, just 120 protein fold-changes are affected by the sub-notch filtering, of which 116 are yeast proteins. The clear majority of fold-changes are further from the ground truth when notch filtering is used (Figure 2d). This observation is counter-intuitive given that sub-notch values are systematically underestimated, suggesting their removal should yield more accurate fold-change estimates. However, since TMT fold-change estimates are always compressed to some extent ^5,7^, the underestimation of these very low intensities below the notch can act to counter the ratio compression. Whilst this is not a justification for retaining the notch to improve fold-change estimates, it further underlines the negligible impact the notch and sub-notch values should have on typical proteomics experiments, where higher level features are typically summarisations of multiple independent PSMs.

An important caveat to our observations is that our benchmark experiment possessed a typical notch prominence and was designed to mimic a simple experiment to detect differential protein abundance. It is possible that in datasets with a more prominent notch, the underestimated tag intensities could become more problematic. Additionally, the notch could conceivably be more problematic where accurate quantification of the ratios between tags are more important than identifying significant differences between treatment groups. For example, in thermal proteome profiling, sub-notch intensities could lead to more poorly fitted curves or models in some edge cases ^13^. If sub-notch values were found to be detrimental in specific applications, we expect that imputing sub-notch values will rectify this. Setting aside these caveats, we believe this artifact in existing Orbitrap Fusion, Lumos and Eclipse quantification data using Tune < 3.0 can be safely ignored for routine proteomics experiments.

## Experimental section

### Cell culture and harvest

*S. cerevisiae* (EOROSCARF; BY4742 (MATα his3Δ1 leu2Δ0 lys2Δ0 ura3Δ0)) were inoculated into YPD media (Bacto-peptone, yeast extract, 50% glucose) at 32°C with constant agitation. Cells were collected by centrifugation when grown to an optical density OD600 ~ 0.5, corresponding to exponential phase, and snap-frozen in liquid nitrogen until lysis. Cell pellets were resuspended in lysis buffer (200 mM HEPES (pH 8.5), 8M urea, 0.2% SDS) supplemented with protease and phosphatase inhibitors (Roche cOmplete mini EDTA-free protease inhibitor cocktail; 11873580001, Roche PhosSTOP; 4906845001) and mechanically disrupted with glass beads (MERCK; G8772) using a FastPrep-24™ 5G (MP Biomedicals SKU; 116005500) with the manufacturer’s pre-defined programme for yeast sample lysis (40 sec mixing at 40 m/s). The lysate was centrifuged to pellet cell debris and supernatant transferred to a fresh tube.

Human epithelial bone osteosarcoma, U-2 OS (ATCC® HTB-96™) cells were cultured and incubated in McCoy’s 5A (Gibco™; 16600082) supplemented with 10% FBS at 37°C in humidified conditions with 5% CO2 and tested to confirm absence of Mycoplasma. Cells were harvested at ~90% confluence by scraping directly from the plate in chilled lysis buffer (as used in yeast lysis) and sonicated on high setting for a total of 15 minutes (30 sec cycles) at 4°C using a Bioruptor® Plus. The lysate was centrifuged to pellet cell debris and supernatant transferred to a fresh tube.

### Reduction, alkylation & digestion

Nuclease enzyme (Millipore Benzonase® Nuclease HC; 71206-3) was added to cell lysates to breakdown interfering DNA before quantification of protein concentration by BCA assay (Thermo Scientific 23225) according to manufacturer instruction. Disulphide bonds in the lysates were reduced with 15 mM dithiothreitol (DTT) for 1 hr at 37°C, followed by alkylation with 55 mM iodoacetamide (IAA) for 1 hr at room temperature in the dark. To remove urea and other substances that could interfere with digestion, lysates were precipitated using chilled 50:50 ethanol:acetone overnight at −20°C. The resulting protein pellets were then resuspended in 200 mM HEPES (pH 8.5) and sonicated for a total of 15 minutes (30 sec cycles) to break up the pellets. Proteins were digested in a two-step digestion with 100:1 protein:protease ratio of Lys-C (Promega; V1671) 37०C for 4 hr, followed by 100:1 trypsin digestion (Promega; V5111) at 37०C overnight.

### Peptide mixing & TMT labelling

Peptide concentrations were measured by using a fluorometric peptide assay (Pierce; 23290) before preparing human/yeast species mixes to specified ratios. 5 μg yeast peptides were used for tags 126, 127N, 127C and 128N, 10 μg for 128C, 129N and 129C, and 30 μg for 130N, 130C and 131. The samples were made up to 100 μg using human peptides, TMT-labelled (Thermo Scientific; 90406) according to manufacturer’s instructions, pooled and lyophilized.

### Peptide clean up & offline pre-fractionation

The multiplexed sample was desalted using solid phase extraction (SPE) with a C18 cartridge (Waters SepPak®; WAT054955) by binding and washing peptides with 0.1% trifluoroacetic acid (TFA) and eluting desalted peptides with 70% acetonitrile/0.05% acetic acid. Peptides were separated using a basic pH reverse-phase liquid chromatography (RP-LC) on a Acquity UPLC system with a diode array detector (210-400 nm) to monitor elution profiles. Peptides were eluted with an Acquity UPLC BEH C18 column (2.1-mm ID × 150-mm; 1.7-μm particle size) (Waters; 186002353) over a 50 min linear gradient from 5 to 75% acetonitrile in ammonium formate (pH 10.0) at a flow rate of 0.244 mL/min. A total of 34 fractions were taken from the elution gradient and concatenated into 15 fractions. Samples were subsequently dried and solubilised in 0.1% formic acid.

### Mass spectrometry data acquisition

TMT labelled samples were analysed using a Dionex Ultimate 3000 RSLC nanoUPLC (Thermo Fisher Scientific Inc, Waltham, MA, USA) system online with an Orbitrap Lumos mass spectrometer (Thermo Fisher Scientific Inc, Waltham, MA, USA), data was collected using both Tune 2.1 and 3.4 version Peptides were loaded onto a trap-column (Thermo Scientific PepMap 100 C18, 5 μm particle size, 100A pore size, 300 μm i.d. x 5mm length) and separation of peptides was performed by C18 reverse-phase chromatography at a flow rate of 300 nL/min and a Thermo Scientific reverse-phase nano Easy-spray column (Thermo Scientific PepMap C18, 2μm particle size, 100A pore size, 75μm i.d. x 50cm length). All samples were acquired in a 120 min run applying data acquisition using synchronous precursor selection MS^3^ (SPS-MS3) ^8^. Analytical chromatography consisted of Buffer A (HPLC H_2_O, 0.1% formic acid) and Buffer B (80% ACN, 0.1% formic acid). 0-3 min at 2% buffer B, 3-93 min linear gradient 2% to 40% buffer B, 93-100 min linear gradient 40% to 90% buffer B, 100-104 min at 90% buffer B, 104-105 min linear gradient 90% to 2% buffer B and 105-120 min at 5% buffer B.

All *m/z* values of eluting peptide ions were measured in an Orbitrap mass analyzer, set at a resolution of 120,000 and were scanned between *m/z* 380-1500 Da. Data dependent MS/MS scans (3 second duty cycle time) were employed to automatically isolate and fragment precursor ions using Collisional-Induced Dissociation (CID) (Normalised Collision Energy of 35%). Only precursors with charge between 2 to 7 were selected for fragmentation, with an AGC target of 10,000 and maximum accumulation time of 50 ms. Precursor isolation was performed by the quadrupole with 0.7 m/z transmission window. MS2 fragments were measured with the Ion Trap analyser. Dynamic exclusion window was set to 70 seconds. SPS ions were all selected within the 400–1,200 m/z range. AGC targets and maximum accumulation times were set to 50,000 and 120 ms respectively. Ten co-selected precursors for SPS-MS3 underwent Higher energy Collisional-induced Dissociation (HCD) fragmentation with 65% normalized collision energy and were analysed in the Orbitrap with nominal resolution of 50 000. Data was acquired with equivalent parameters for both version of Tune, 2.1 and 3.4.

### Mass spectrometry data analysis

Raw data were viewed in Xcalibur v3.0.63 (Thermo Fisher Scientific), and data processing was performed in Proteome Discovered v2.4 (Thermo Fischer Scientific). Reference *Homo sapiens* and *Saccharomyces cerevisiae* fasta databases containing all review UniProt/Swiss-Prot entries were downloaded from www.uniprot.org on April 2018 and June 2020, respectively. The raw files were submitted to a database search using PD with Sequest HF algorithm using the concatenated reference databases and the Common contaminant database ^14^. The peptide and fragment mass tolerances were set to 10 ppm and 0.5 Da, respectively. Static modification carbamidomethyl on cysteine was applied as well as TMT-6plex tagging of lysines and peptide N terminus. Oxidation of methionine and deamidation of asparagine and glutamine were included as variable modifications and up to two missed cleavages were allowed. Percolator node was used for false discovery rate estimation and only rank one peptides identifications of high confidence (FDR < 1%) were accepted.

Previously published data ^4,11^ was downloaded from pride accessions PXD011254 and PXD007683 in raw format. Raw data was re-analysed as indicated above, except that the reference proteomes for *H.sapiens* (UP000005640) and *Saccharomyces cerevisiae* (UP000002311) were downloaded in fasta format on 17 January 2020 and used for database searching with Mascot server (ver. 2.4, Matrix Science)

### Data processing and analysis

PSM-level quantification was exported from Proteome Discoverer in text format. Downstream processing, analysis, aggregation and visualisation performed using R ^15^ v4.0.3 “Bunny-Wunnies Freak Out” and the tidyverse ^16^, MSnbase ^17^ and camprotR (https://github.com/CambridgeCentreForProteomics/camprotR) packages.

PSM quantifications were filtered to remove potential contaminating proteins using the cRAP database ^14^. Based on a visual assessment of the tag intensity distributions, the approximate boundaries of the notch were denoted as 4.25 and 5.75.

For the prediction of expected reporter ion intensities with the reanalysed data, PSMs with the same sequence within an experiment were grouped, excluding groups with only one PSM, or where the highest intensity PSM contained missing values. Intra-group intensity adjustment factors were then calculated, representing the intensities of each PSM relative to the most intense PSM, using only the channels without missing values. The highest intensity PSM was then normalised by dividing each tag intensity by the total PSM intensity, such that the total normalised intensity was 1. These normalised intensities were then multiplied by the adjustment factors to yield tag intensity predictions for each PSM in the group, except the highest intensity PSM.

To explore the relationship between tag intensities in the benchmark dataset, individual tag intensities were compared to the mean value for a comparator group of tags. A generalized additive smoothing model for the relationship between tag intensity and intensity difference was fitted using the default model for the ggplot function geom_smooth with method=’gam’, namely the gam function in mgcv, with options ‘formula = y ~ s(x, bs = “cs”)’ and ‘method = “REML”’.

Prior to PSM filtering, to remedy any small difference in the total tag intensity in each channel, PSM intensities were log center-median normalised, before exponentiating back to the untransformed values.

PSM filtering was performed using 4 PSM metrics: (1) Delta score, which represents the normalised difference in the spectrum matching scores between the top ranked and second ranked peptides, calculated as delta = (rank 1 score - rank 2 score) / rank 1 score, and is thus bounded between 0 (no difference) and 1 (no rank 2 score). (2) Co-isolation, calculated as: 100 * (1 - (precursor intensity in isolation window / total intensity in isolation window)). (3) Average reporter signal/noise, representing the average tag intensity for the PSM. (4) Presence of values within or below the notch, where the upper boundary of the notch was used to identify PSMs with any values below this threshold. The following values were used to filtering PSMs according to these metrics: (1) Delta score > 0, 0.2 or 0.5, (2) Co-isolation < 100 %, 50 % or 10 %, (3) Average reporter S/N > 0 or 10 (4) Notch PSMs retained or removed. All combinations of filtering thresholds were performed, yielding 36 sets of filtered PSMs.

Aggregation to protein-level intensities involved removal of PSMs with missing values, removal of proteins without at least two PSMs, and summation of PSM-level tag intensities. Aggregation of PSM intensities to peptide-level intensities proceeded in the same manner, but demanding at least two PSMs per peptide sequence.

### Statistical analysis

To identify peptides and proteins with significant differential abundance, we used *limma* ^18^ *Limma* was run with default settings, with the exception that we set trend=TRUE for the eBayes function call, so that the prior variance was dependent on the trend between feature intensity and observed variance. Protein intensity was modeled to depend on condition, with results extracted for the contrast between 1x vs 2x and 1x vs 6x tags. P values were adjusted for multiple testing within each PSM filtering schema, using the Benjamini-Hochberg False Discovery Rate (FDR) procedure ^19^. Features with FDR < 1 % were deemed to have significantly different intensity.

Yeast proteins with a significant increase in intensity and human proteins with a significant decrease in intensity were deemed true positives (TP). Vis versa, significant changes in the opposite direction were deemed false positives (FP). Proteins without significant changes in intensity were deemed false negatives (FN).

F1 scores were calculated as the harmonic mean of recall (TP / (TP + FN)) and precision (TP / (TP + FP)). No proteins were deemed true negatives (TN) since all proteins should have a change in intensity.

### Extracting tune version from PRIDE submissions

Dataset identifiers for 7072 studies using Orbitrap were obtained from ProteomeCentral (http://proteomecentral.proteomexchange.org/cgi/GetDataset) by searching for entries with ‘Orbitrap’ in the instrument text on 28 June 2021. These were further narrowed down to 1525 studies in the PRIDE repository with ‘Orbitrap Fusion’, ‘Orbitrap Fusion Lumos’ or ‘Orbitrap Eclipse’ listed as instrument since these share the same series of signal acquisition software. Studies with multiple instruments listed were ignored to avoid incorrectly asserting the version of Tune used. For each study, the smallest .raw file was downloaded and the meta information extracted using the R package rawDiag ^20^, including the version of the aquisition software, Tune. For 242 / 1525 studies, the Tune version was not extracted either because the instrument detailed in the .raw file was not one of the above Orbitraps, no .raw files were found, the downloaded .raw file could not be parsed by rawDiag, or the .raw file URLs did not permit downloading.

## Data availability

The mass spectrometry proteomics data have been deposited to the PRIDE Archive (http://www.ebi.ac.uk/pride/archive/) via the PRIDE partner repository with the data set identifier PXD027248.

All analyses and results are publically available from the github repo https://github.com/CambridgeCentreForProteomics/notch, alongside the peptide spectrum match (PSM)-level quantification from Proteome Discoverer, and reference fasta databases used. Release v0.1 of the repository used for this manuscript is archived with DOI 10.5281/zenodo.5105723.

## Acknowledgements

We would like to thank Nianshu Zhang for kindly providing *S.cerevisiae* cultures, Mie Monti for kindly preparing *S.cerevisiae* cell lysates and Rayner Queiroz for kindly providing U-2 OS cultures. We would also like to thank Mike Deery and members of the Lilley group for insightful discussions regarding the source and impact of the notch.

## Author Contributions

**TSS**: Conceptualization, Methodology, Formal analysis, Data Curation, Writing - Original Draft, Writing - Review & Editing, Visualization, Project administration. **AA**: Methodology, Investigation, Writing - Review & Editing. **JAC**: Methodology, Investigation,Writing - Review & Editing. **OMC**: Methodology, Writing - Review & Editing, Funding acquisition. **ME**: Formal analysis, Writing - Review & Editing, **KSL**: Methodology, Writing - Review & Editing, Supervision, Funding acquisition.

## Supplementary figures

**Figure S1.**
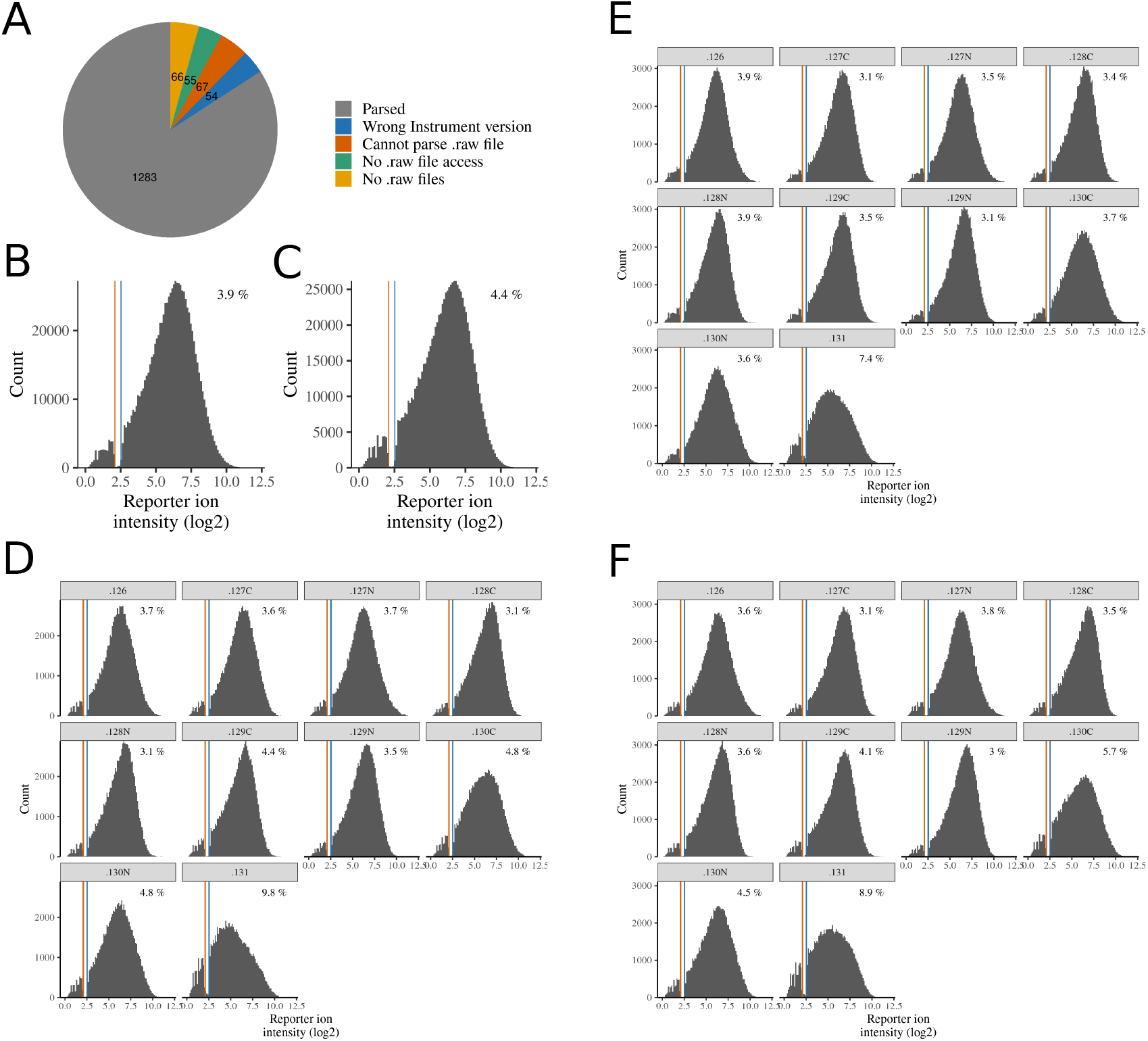
(A) The Tune version could be ‘Parsed’ from 1283 studies, with the remaining not accessible for the reasons described. (B-C) The distribution of ion signals for TMT reporters for U-2 OS LOPIT-DC replicate 2 (B) and replicate 3 (C). The approximate boundaries of the notch region are denoted by vertical lines. The percentage of tag intensities below the upper boundary of the notch is stated in the top right corner. (D-F) The distribution of ion signals for each individual TMT reporter; replicate 1 (D), replicate 2 (E) and replicate 3 (F).

**Figure S2:**
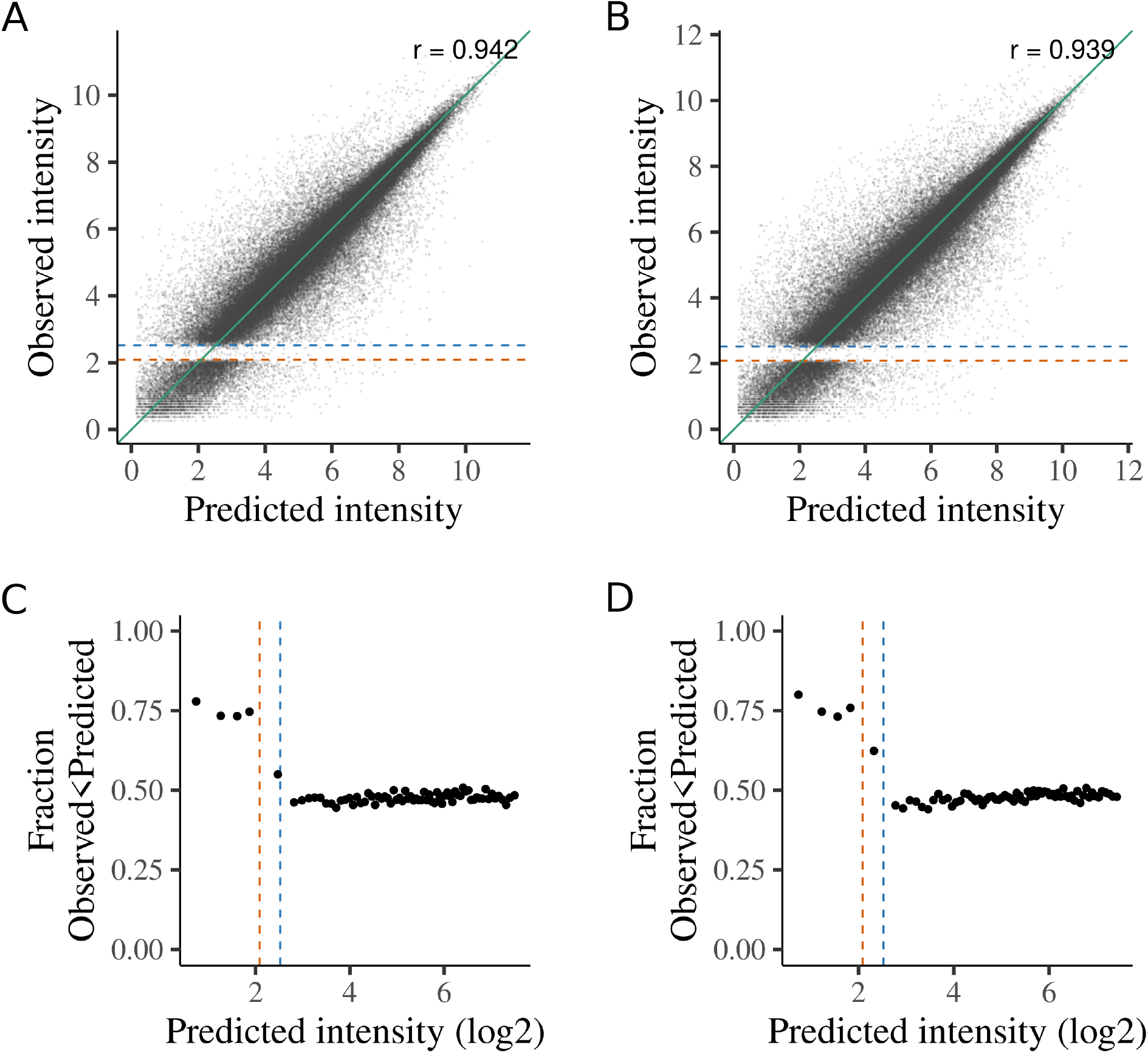
Characterisation of the reporter ion intensities around the notch in U-2 OS LOPIT-DC (A-B) Observed tag intensities vs predicted tag intensities for replicate 2 (A) and replicate 3 (B). Green line is equality. Pearson product-moment correlation coefficient shown in the top right corner. (C-D) Fraction of observed reporter ion intensities that are below the prediction for replicate 2 (C) and replicate 3 (D). Observed tag intensities below the notch are systematically underestimates relative to the prediction.

**Figure S3.**
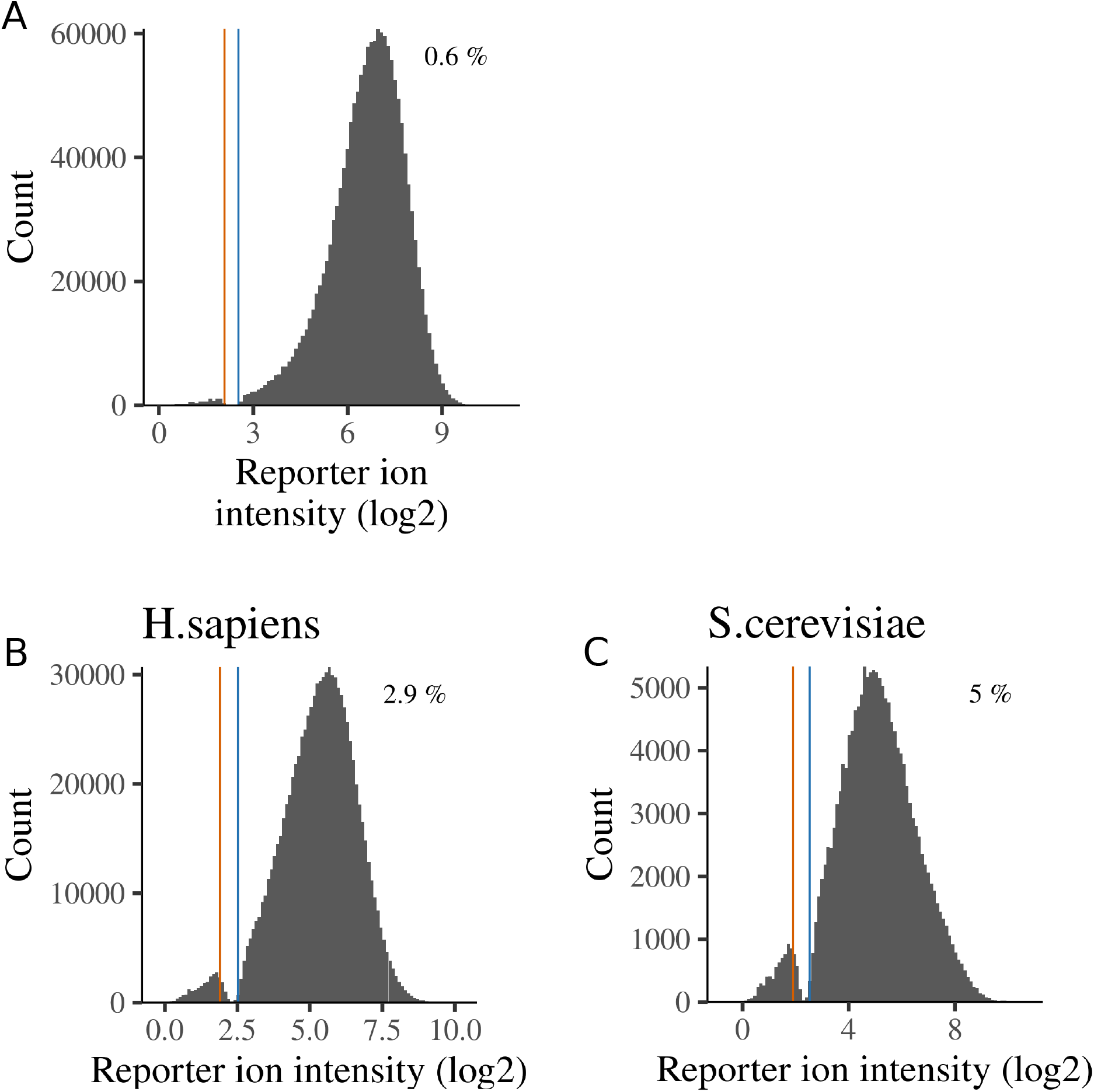
Distributions for reporter tag intensities in O’connell et al data (A; ^4^) and the benchmark dataset, considering only human (B) or yeast (C) proteins. The percentage of tag intensities below the upper boundary of the notch is stated in the top right corner.

**Figure S4.**
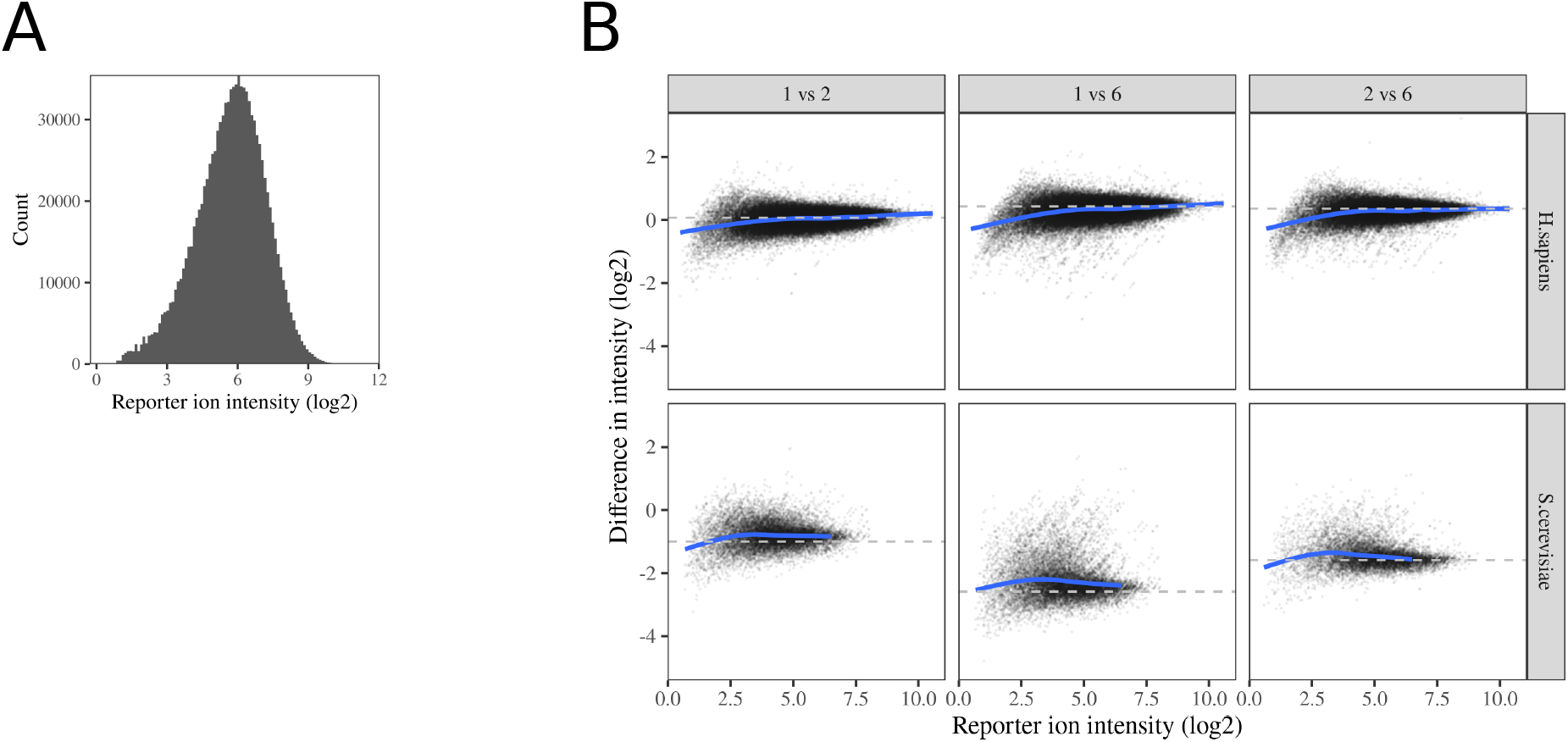
Reporter ion intensity and fold-changes for benchmark dataset when using Tune 3.4.(A) The distribution of reporter ion intensities for TMT reporters. No notch is observed. (B) The difference between a single tag intensity and the mean tag intensity for a comparator group of tags. The ground truth is denoted by a dashed horizontal line. The blue line presents a generalized additive smoothing model for the relationship between tag intensity and intensity difference.

**Figure S5.**
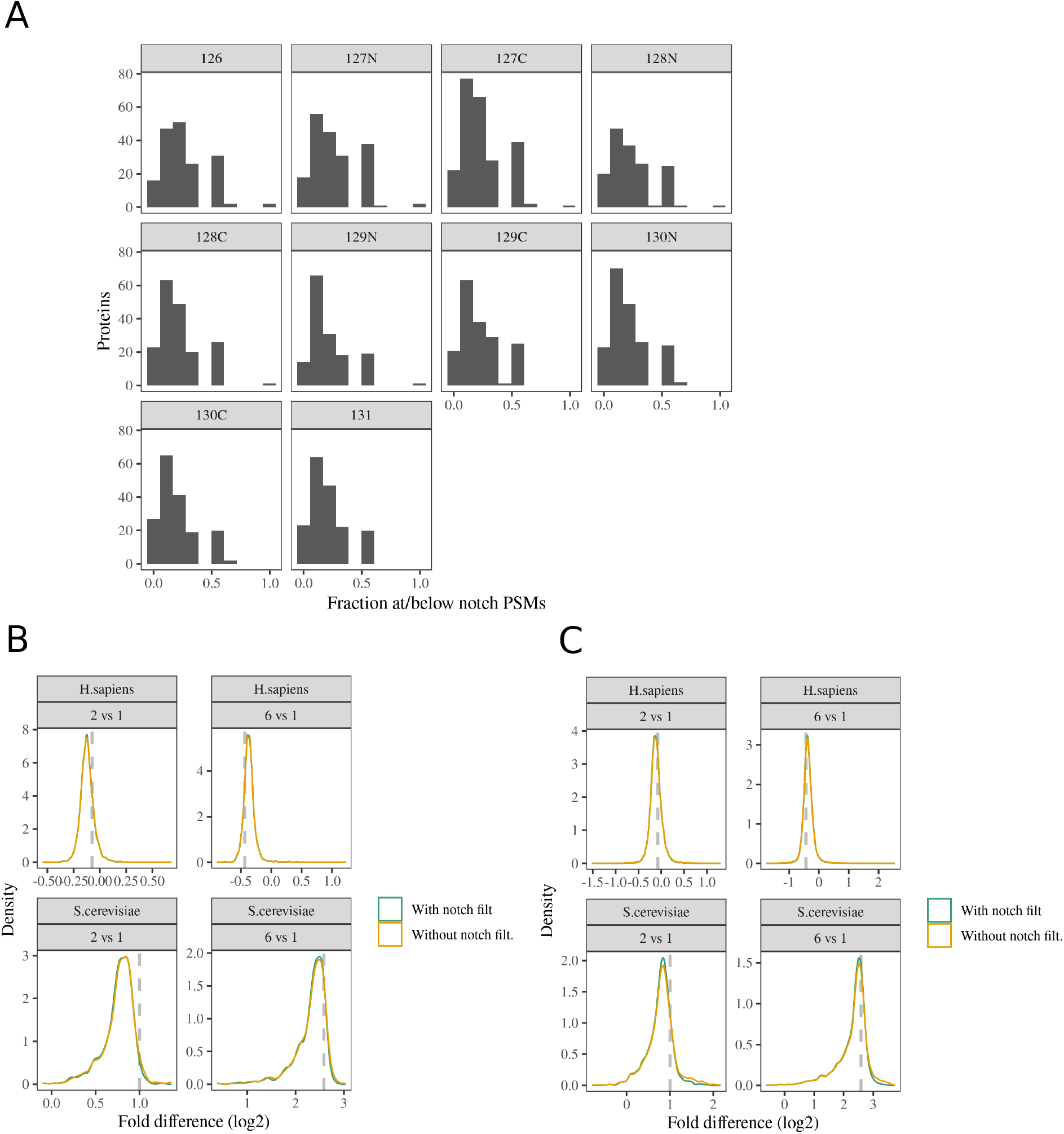
(A) Tallies for the fraction of the aggregated reporter ion intensities at or below the notch for each protein. Proteins with no intensities at or below notch are not tallied. (B-C) Observed fold-changes between tag groups, with and without notch filtering, for proteins (B) and peptides (C).

**Figure S6.**
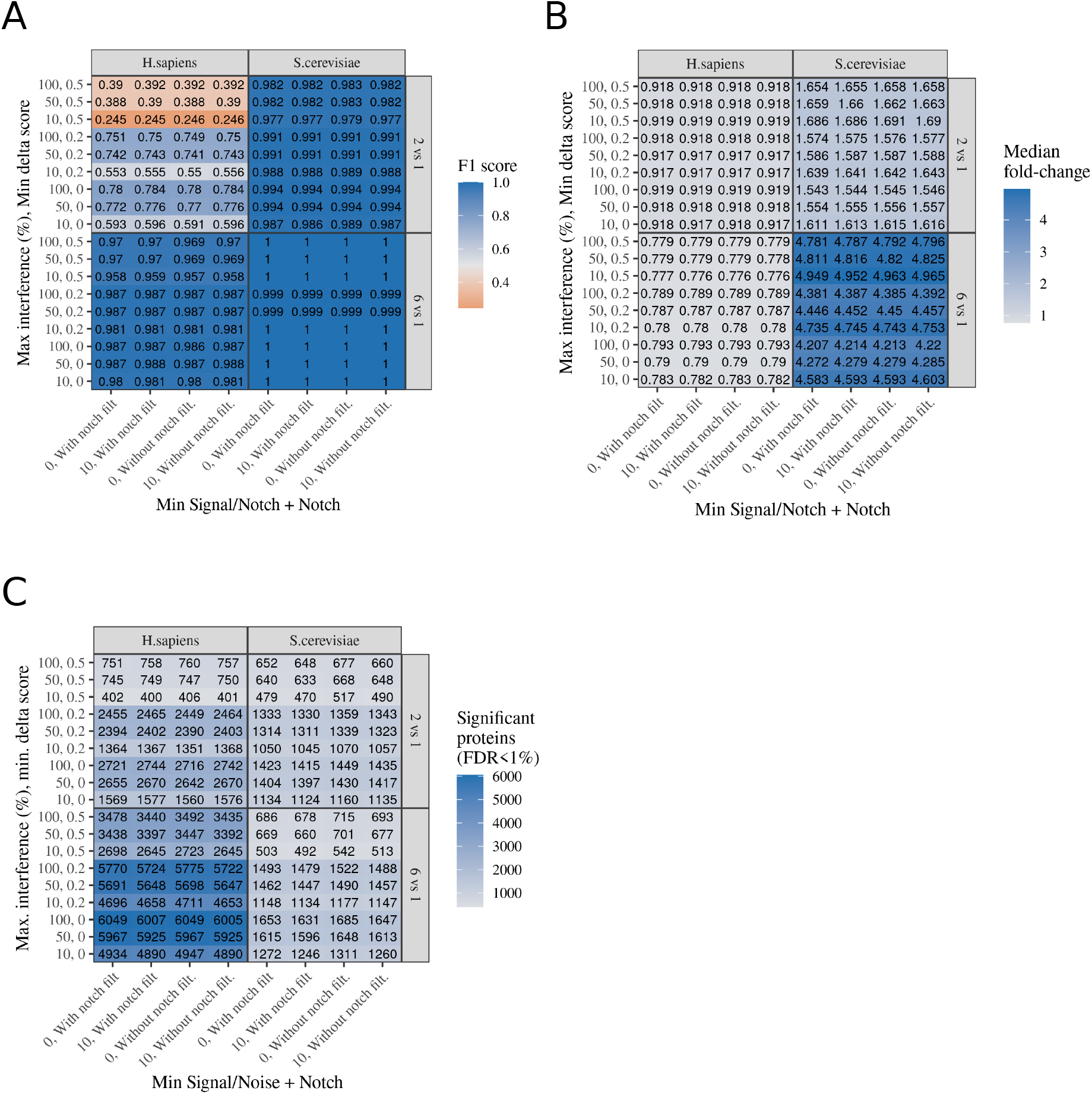
Summary of limma analysis of protein-level intensities. Limma analysis was performed on datasets generated from multiple different PSM filtering schemas, as indicated on the x-axis and y-axis. Each axes describe the combination of two filtering parameters. ‘Min Signa/Noise’ is the minimum average signal/noise for the PSM. ‘+ Notch’ describes whether PSMs containing any intensities at or below the notch were removed. ‘Max interference (%)’ describes the maximum allowed interference/co-isolation. ‘Min delta score’ describes the minimum required delta score. (A) Accuracy of differential protein intensity detection, presented as the F1 score (harmonic mean of precision and recall). Only proteins present with all PSM filtering schemas were included (B) Median fold change between groups of tags. Only proteins present with all PSM filtering schemas were included. (C) The number of proteins with significantly different intensity.

**Figure S7.**
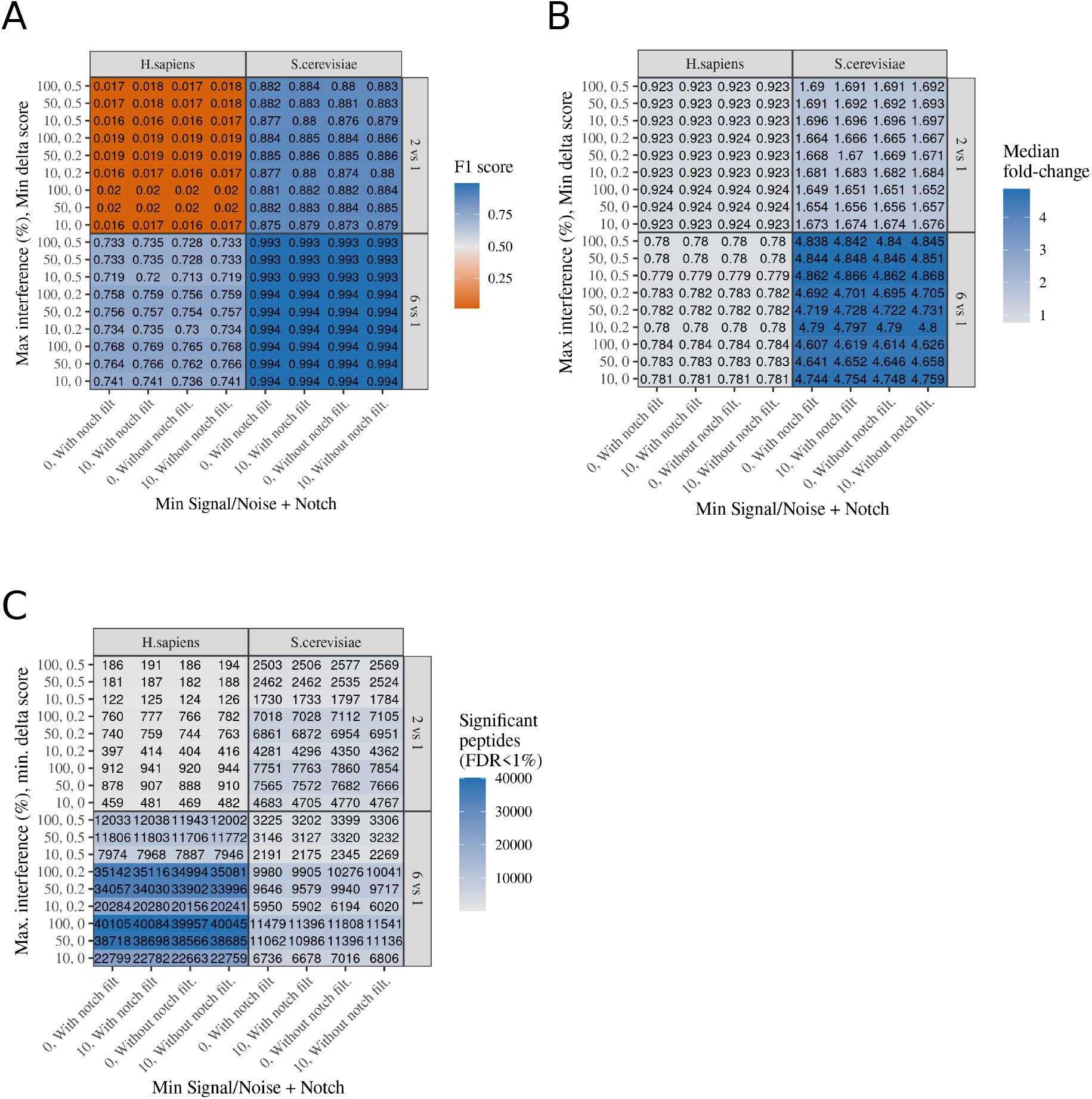
Summary of limma analysis of peptide-level intensities. Limma analysis was performed on datasets generated from multiple different PSM filtering schemas, as indicated on the x-axis and y-axis. Each axes describe the combination of two filtering parameters. ‘Min Signa/Noise’ is the minimum average signal/noise for the PSM. ‘+ Notch’ describes whether PSMs containing any intensities at or below the notch were removed. ‘Max interference (%)’ describes the maximum allowed interference/co-isolation. ‘Min delta score’ describes the minimum required delta score. (A) Accuracy of differential protein intensity detection, presented as the F1 score (harmonic mean of precision and recall). Only peptide present with all PSM filtering schemas were included (B) Median fold change between groups of tags. Only peptide present with all PSM filtering schemas were included. (C) The number of peptides with significantly different intensity.

